# Increasing consensus of context-specific metabolic models by integrating data-inferred cell functions

**DOI:** 10.1101/384099

**Authors:** Anne Richelle, Austin W.T. Chiang, Chih-Chung Kuo, Nathan E. Lewis

**Author notes:** Correspondence (N.E.L).

## Abstract

Genome-scale metabolic models provide a valuable context for analyzing data from diverse high-throughput experimental techniques. Models can quantify the activities of diverse pathways and cellular functions. Since some metabolic reactions are only catalyzed in specific environments, several algorithms exist that build context-specific models. However, these methods make differing assumptions that influence the content and associated predictive capacity of resulting models, such that model content varies more due to methods used than cell types. Here we overcome this problem with a novel framework for inferring the metabolic functions of a cell before model construction. For this, we curated a list of metabolic tasks and developed a framework to infer the activity of these functionalities from transcriptomic data. We protected the data-inferred tasks during the implementation of diverse context-specific model extraction algorithms for 44 cancer cell lines. We show that the protection of data-inferred metabolic tasks decreases the variability of models across extraction methods. Furthermore, resulting models better capture the actual biological variability across cell lines. This study highlights the potential of using biological knowledge, inferred from omics data, to obtain a better consensus between existing extraction algorithms. It further provides guidelines for the development of the next-generation of data contextualization methods.

## Introduction

Genome-scale metabolic models (GeMs) have been widely used for model-guided analysis of large-scale omics datasets, since they provide cellular context to these data by establishing a mechanistic link from genotype to phenotype. GeMs include all reactions in an organism. Since not all enzymes are active in each cell type or culture condition, algorithms have been developed to build context-specific models using omics data to recapitulate the metabolism of specific cell types under specific conditions (Opdam *et al*, 2017; Robaina Estévez & Nikoloski, 2014). Each of these algorithms have provided useful insights in the metabolism of specific cell and tissue types (Opdam *et al*, 2017; Wang *et al*, 2012b; Zur *et al*, 2010; Vlassis *et al*, 2013; Becker & Palsson, 2008; Agren *et al*, 2012; Jerby *et al*, 2010; Hefzi *et al*, 2016; Kumar *et al*, 2014). However, since each of these methods use different assumptions to guide reaction inclusion and removal, they result in considerable differences in size, functionality, accuracy, and ultimate biological interpretation, even when using the same data set (Opdam *et al*, 2017; Ferreira *et al*, 2017; Robaina Estévez & Nikoloski, 2014).

The poor consensus in generated models requires increased caution in the interpretation of model-derived hypotheses of how metabolism is used under specific environments. Indeed, most generated models, upon construction, will be missing known metabolic functions and this varies considerably for models built using different approaches (Opdam *et al*, 2017). To gain confidence in model predictions and reconcile the differences across approaches, users can enforce the inclusion of known metabolic capabilities in the model. In this regard, the tINIT extraction algorithm introduced the possibility to enforce the capacity of context-specific models to represent some cellular functionalities by using a list of metabolic tasks known to occur in all cell types (Agren *et al*, 2014). However, this protectionist approach requires one to know and predefine the functionalities of a specific cell line, tissue, or context.

To overcome this, we propose an approach to infer the functionalities of a cell or tissue from omics data, and then protect these functions to guide the construction of a context-specific model. To this end, we curated and standardized published lists of metabolic tasks (Blais *et al*, 2017; Thiele *et al*, 2013), resulting in a collection of 210 tasks covering 7 major metabolic activities of a cell (energy generation, nucleotide, carbohydrates, amino acid, lipid, vitamin & cofactor and glycan metabolism). We also developed a framework to directly predict the activity of these functionalities from transcriptomic data and subsequently use these for a protectionist approach to diverse existing extraction algorithms. Models resulting from this approach should more comprehensively capture the unique metabolic functions of a given cell type. We further evaluated the validity and variation across models built with this approach, coupled to existing context-specific extraction methods. Specifically, we constructed hundreds of models for 44 cancer cell lines in which we built the models using standard approaches or protected a list of metabolic functions that have been inferred from the original transcriptomic data of each cell line. We also varied the reference human reconstruction and algorithms employed for the generation of cell line specific models, using two different reference models (iHsa (Blais *et al*, 2017) and Recon2.2 (Swainston *et al*, 2016b)) and 6 different algorithms (mCADRE (Wang *et al*, 2012a), fastCORE (Vlassis *et al*, 2013), GIMME (Becker & Palsson, 2008), INIT (Agren *et al*, 2012), iMAT (Zur *et al*, 2010), and MBA (Jerby *et al*, 2010)). We compared the sets of extracted models at the level of reaction content, metabolic functions, and capacity to predict gene essentialities using CRISPR-Cas9 loss-of-function screens. Through this study, we highlight the value of using experimental data to help infer the set of metabolic tasks that should be included in a model, in an effort to obtain greater consensus across existing extraction algorithms.

## Results

### Context-specific extraction methods yield more variation in model content than omics data of different cell lines

We built models from Recon 2.2 (Swainston *et al*, 2016a) and iHsa (Blais *et al*, 2017) using six model extraction methods (MEMs: mCADRE, fastCORE, GIMME, INIT, iMAT, MBA) for 44 different cell lines from the NCI-60 panel (Supplementary Table 1; 15 cell lines were not used due to the absence RNA-Seq data in CellMiner for these cell lines (Reinhold *et al*, 2012)). Uptake and secretion rates of the input GeMs were quantitatively constrained using a list of experimentally measured metabolites (Supplementary Table 2; Jain et al., 2012; Zielinski et al., 2017). Furthermore, a biomass function, consisting of 56 metabolites required for growth, was added and constrained to the experimentally measured growth rate of the cell lines (Supplementary Table 3). The biomass function and constraints from exometabolomic data introduced in the GeMs were implemented as described in Opdam et al. (2017). The extraction process of cell line specific models was done based on RNA-Seq data (Reinhold *et al*, 2012) to specify active genes in each cell line. Details on the implementation of MEMs tested and the preprocessing of gene expression data for the definition of gene activity are provided in the Methods section.

To assess the relative impact of algorithm and data source on model content, we conducted a principal component analysis (PCA) of the reactions in all models for each reference GeM. As observed previously (Opdam *et al*, 2017; Ferreira *et al*, 2017), the decisions regarding algorithm choice significantly impact the content of our cell line-specific models. The first principal (PC1) component explains 38% of the overall variance in model reaction content, with >60% of the variation in PC1 explained by the choice of model extraction method (Figure 1A and 1B). Indeed, the different algorithms yielded cell line-specific models that varied considerably in size, with few reactions common to all models extracted from either Recon2.2 or iHsa (Figure 1C, Supplementary Figure 1). Even among models extracted using the same algorithm, there is non-negligible variability in model reaction content (Figure 1D). This leads to the generation of models that are substantially different with respect to the cell line considered, while the transcriptomic data used to tailor the GeMs shows high consistency across most cell lines (Figure 1E).

**Figurue 1.**
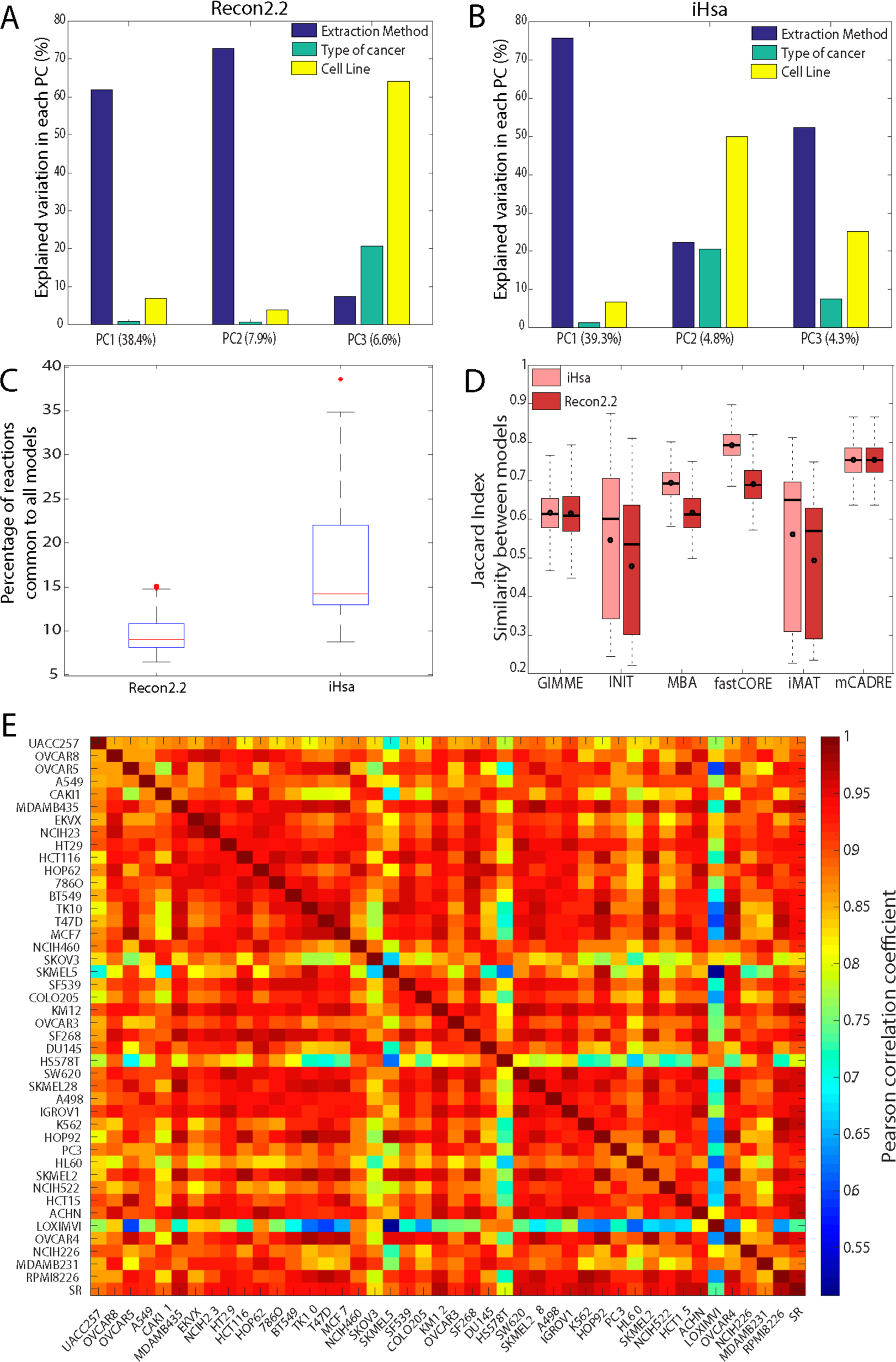
Choice of extraction method is mainly responsible for the variability in the reaction content across models. The extraction method used contributes the most to the first PC for models built using (A) Recon2.2 or (B) iHsa as a reference GeM. (C) Only a small percentage of reactions are shared in all the models extracted from both Recon2.2 (shared reactions = 218) and iHsa (shared reactions = 513). (D) For each method, the similarity of models of different cell lines (computed using a Jaccard index based on the reaction content) varies substantially, while (E) the transcriptomic data used to generate these models present a much higher correlation between cell lines.

### Metabolic tasks as a tool for model benchmarking and model extraction

Model reaction content is often evaluated to compare context-specific algorithms. Recently, approaches to benchmark models with their functionalities have been proposed (Opdam *et al*, 2017; Agren *et al*, 2014; Pacheco *et al*, 2016). Current approaches use repositories of known cellular tasks to assess the capacity of models to achieve specific modeling goals or to enable the representation of specific metabolic functions. This idea of assessing the quality of a metabolic network reconstruction using biological knowledge was introduced in Recon1 through the characterization of the “human metabolic knowledge landscape” (Duarte *et al*, 2007). However, the concept of “metabolic tasks” (Figure 2A) was clearly defined in 2013 by Thiele and coworkers to benchmark the improvements of Recon2 compared to Recon1, wherein they stated that “a metabolic task is defined as a nonzero flux through a reaction or through a pathway leading to the production of a metabolite B from a metabolite A”. Since then, additional lists of tasks have been published. To standardize these and develop a framework for their easy use with GeMs, we curated the existing lists of metabolic tasks (Thiele *et al*, 2013; Blais *et al*, 2017) and obtained a collection of 210 tasks covering 7 major metabolic activities of a cell (energy generation, nucleotide, carbohydrates, amino acid, lipid, vitamin & cofactor and glycan metabolism) (Figure 2B and 2C, Supplementary Table 4). We evaluated the task collection using genome-scale metabolic models for human (Swainston *et al*, 2016a; Thiele *et al*, 2013; Duarte *et al*, 2007; Quek *et al*, 2014; Blais *et al*, 2017), CHO cells (Hefzi *et al*, 2016), rat (Blais *et al*, 2017) and mouse (Sigurdsson et al., 2010; Figure 2D, Supplementary Table 5). Details on our proposed formalism of the metabolic tasks and the associated computation framework for their use are presented in the Methods.

**Figurue 2.**
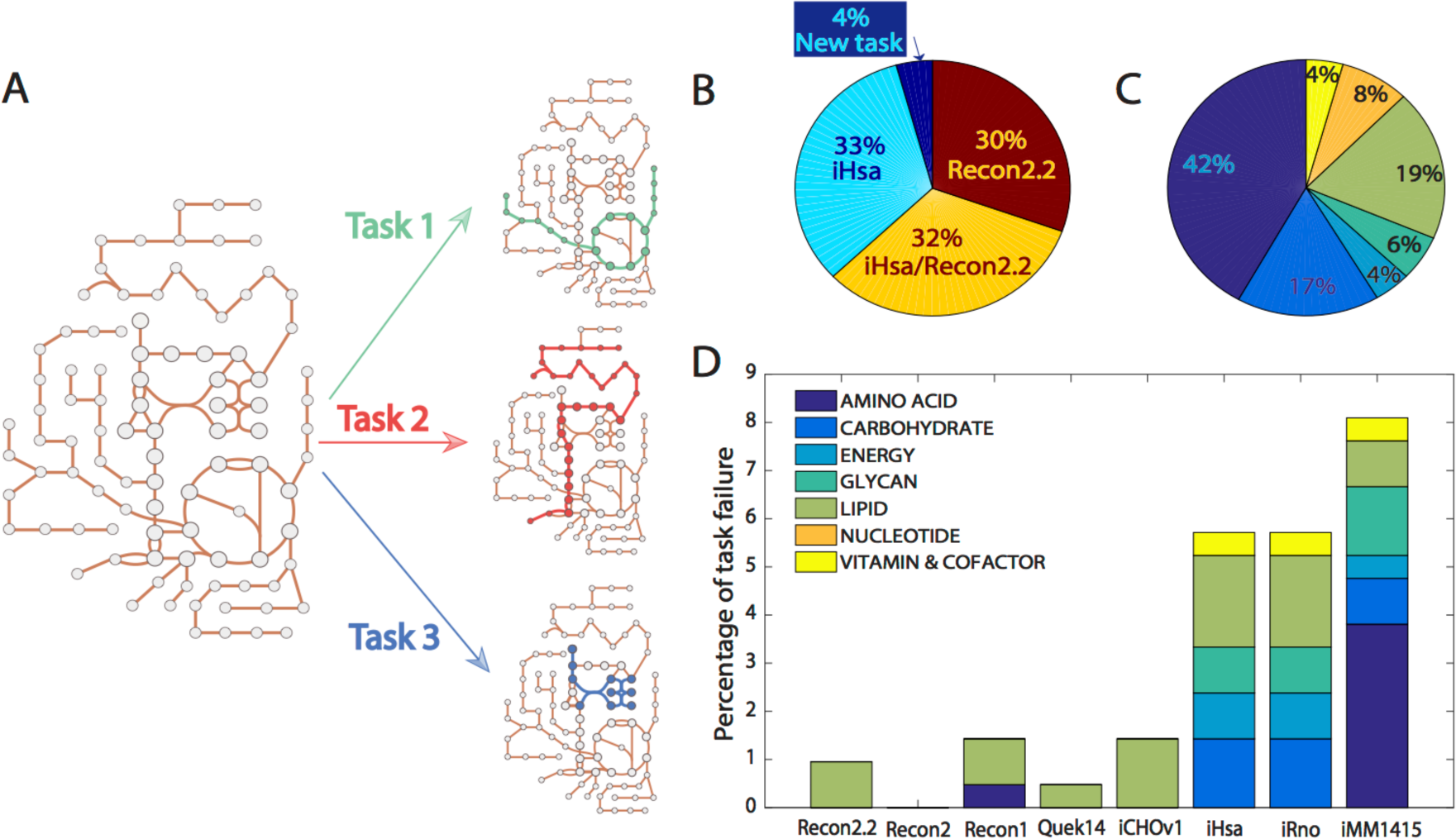
Generation of a collection of 210 tasks representing known metabolic features. (A) A metabolic task can be defined as the set of reactions needed to transform input metabolites into defined products. (B) Original sources of our collection of 210 curated tasks. (C) The curated list of tasks covers 7 main metabolic systems. (D) Several existing genome-scale models were used to evaluate the collection of curated tasks. A small number of tasks were non-functional on specific GeMs, for reasons detailed in Supplementary Table 5.

Metabolic tasks can be used to compare the performance of models extracted from different reference GeMs. As observed at the level of the reaction content, the extraction method strongly influences the model functions (explaining >50% of the overall variance in the first PC; Figure 3A). However, the reference model is the most prominent factor in the second PC underlying a non-negligible influence of this variable in the extraction process. This is mainly due to differences in gene, protein, reaction association (GPR) annotations and reaction content between Recon2.2 and iHsa. Interestingly, Recon2.2 captures more metabolic functions with fewer reactions (Figure 3B). However, the number of successful tasks increases proportionally with the number of reactions in a model. Furthermore, as the extraction method used influences the number of reactions removed, distinct patterns are seen from the ratio of the number of metabolic tasks to the number of reactions introduced by the different algorithms (Figure 3C). As for the reaction content, the number of tasks retained in each model varies substantially, depending on the cell lines considered. Surprisingly, only 8% of the tasks are present in all models (Figure 3D, Supplementary Figure 2), thus highlighting the large variation in metabolic functions a model will have, depending on algorithm choice.

**Figurue 3.**
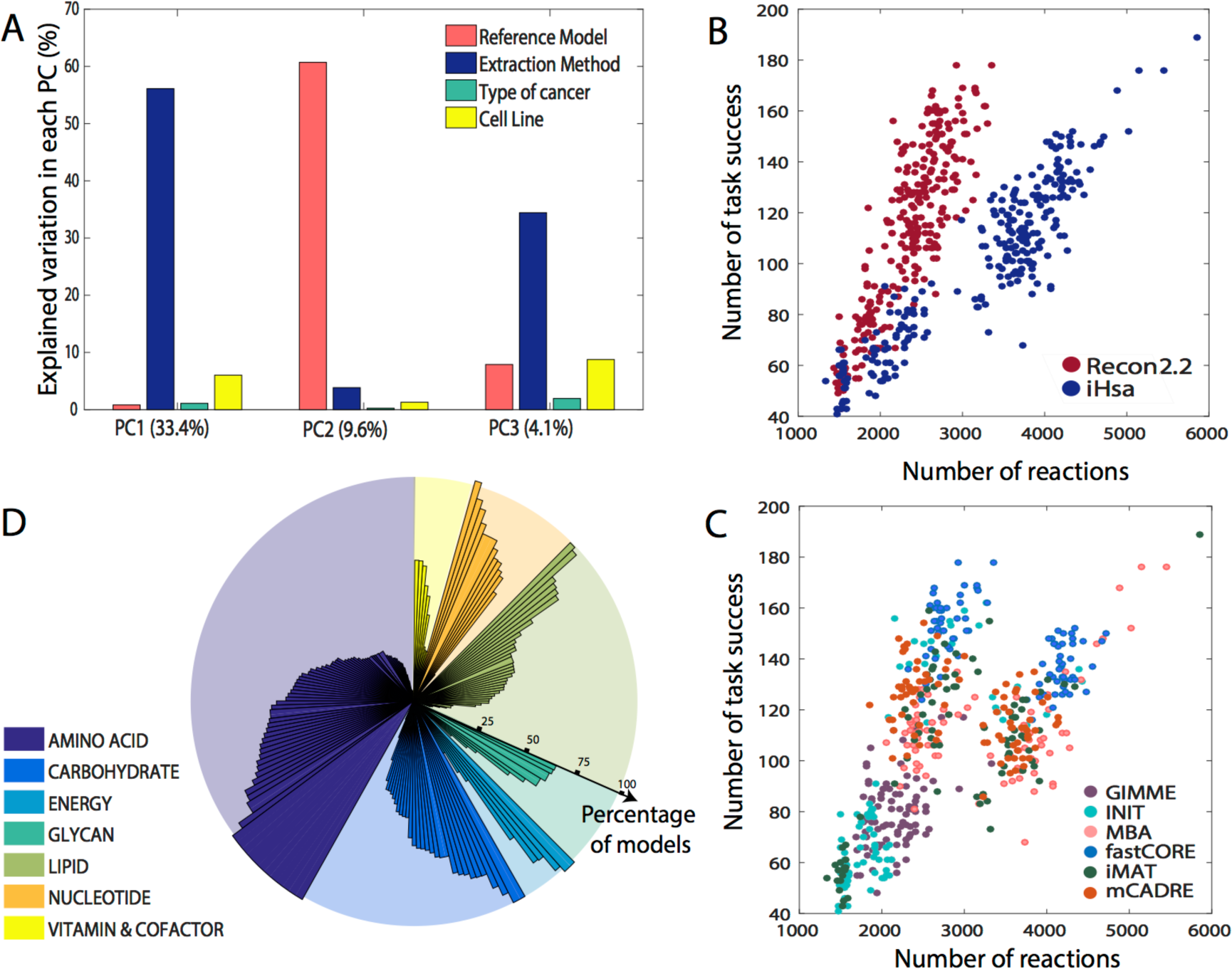
Metabolic tasks can be used to benchmark models. (A) The extraction method used contributes the most to the first PC of functional metabolic tasks and the reference model explains the most variance within the second PC. (B) Recon2.2 captures more metabolic functions with fewer reactions. (C) Some extraction methods are associated with a higher capacity to conserve more metabolic functions, as they typically retain more reactions. (D) The percentage of tasks present in all extracted models is low, and predominantly associated with amino acid metabolism. Furthermore, some metabolic functionalities are not retained in any extracted models.

### Protecting inferred metabolic tasks reduces the variability of model content from different algorithms

We inferred active metabolic tasks directly from transcriptomic data using the whole genome-scale model. To this end, we computed the list of reactions associated with each task and used the GPR rules to determine the gene expression levels associated to each of these reactions. A metabolic score is attributed to each task by using the mean activity level of each reaction (Figure 4A; See Methods). We found that more than the half of the tasks should be conserved across all cell lines (Figure 4B), which is far more than those active using the algorithms in their standard format (i.e., without protecting tasks). Therefore, we generated a new set of models, wherein we also enforced the inclusion of reactions associated with tasks inferred for each of the 44 different cell lines (Supplementary Table 6). We focused on MBA-like algorithms (i.e., MBA, fastCORE and mCADRE), since they are directly amenable to use the protectionist approach with minor modifications to the algorithms (Figure 4C). Indeed, other algorithms do not ensure the inclusion of a reaction even if it is enforced. For example, iMAT relies on the definition of a core set of high-confidence reactions, but core reactions can be removed if it depends on many non-expressed non-core reactions (Figure 4C). See Methods for a detailed description of the implementation of the protectionist approach for each algorithm.

**Figurue 4.**
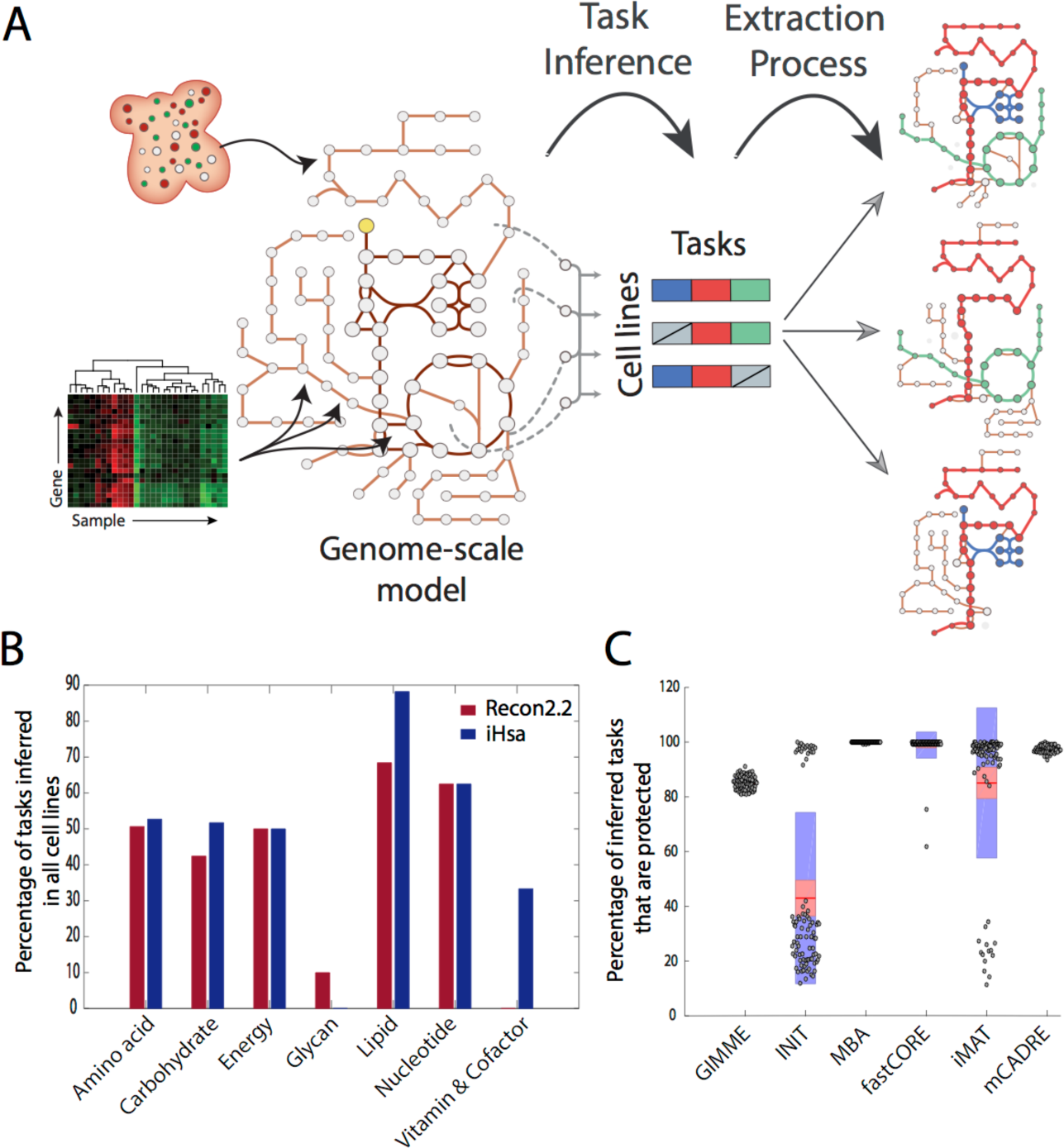
Metabolic tasks can be inferred from omics data to determine which tasks should be protected during the model extraction process. (A) Metabolic functions are inferred from transcriptomic data using the genome-scale model and then protected during the implementation of the extraction algorithms. (B) The functional analysis of transcriptomic data highlights that more than half of the tasks should be conserved across all the cell lines. (C) The computational framework of some extraction algorithms does not allow a complete protection of the inferred metabolic tasks, but protected tasks were almost completely retained for MBA-like methods (i.e. MBA, fastCORE and mCADRE).

For equivalent extraction setups (i.e., same reference model, extraction method, and cell line), the number of reactions included in the extracted model was not considerably influenced by the protection of the metabolic task, while the number of active tasks clearly increases (Supplementary Figure 3). We performed PCA of the reaction content and the metabolic functions of the models with protected tasks. We observed that the protection of metabolic tasks inferred from data significantly decreased the influence of the extraction method on the final model content. However, the use of this approach remains sensitive to the choice of the reference model (Figures 5A-B). Furthermore, we observed that the variation in model content was better explained by the cell lines (Supplementary Figure 4). Actually, the task protection increases the similarities between context-specific models with respect to the cell line (Supplementary Figure 5) but also with respect to the transcriptomic data (Supplementary Figure 6). Finally, all the models now share more than 64% of the metabolic tasks (Figure 5C).

**Figurue 5.**
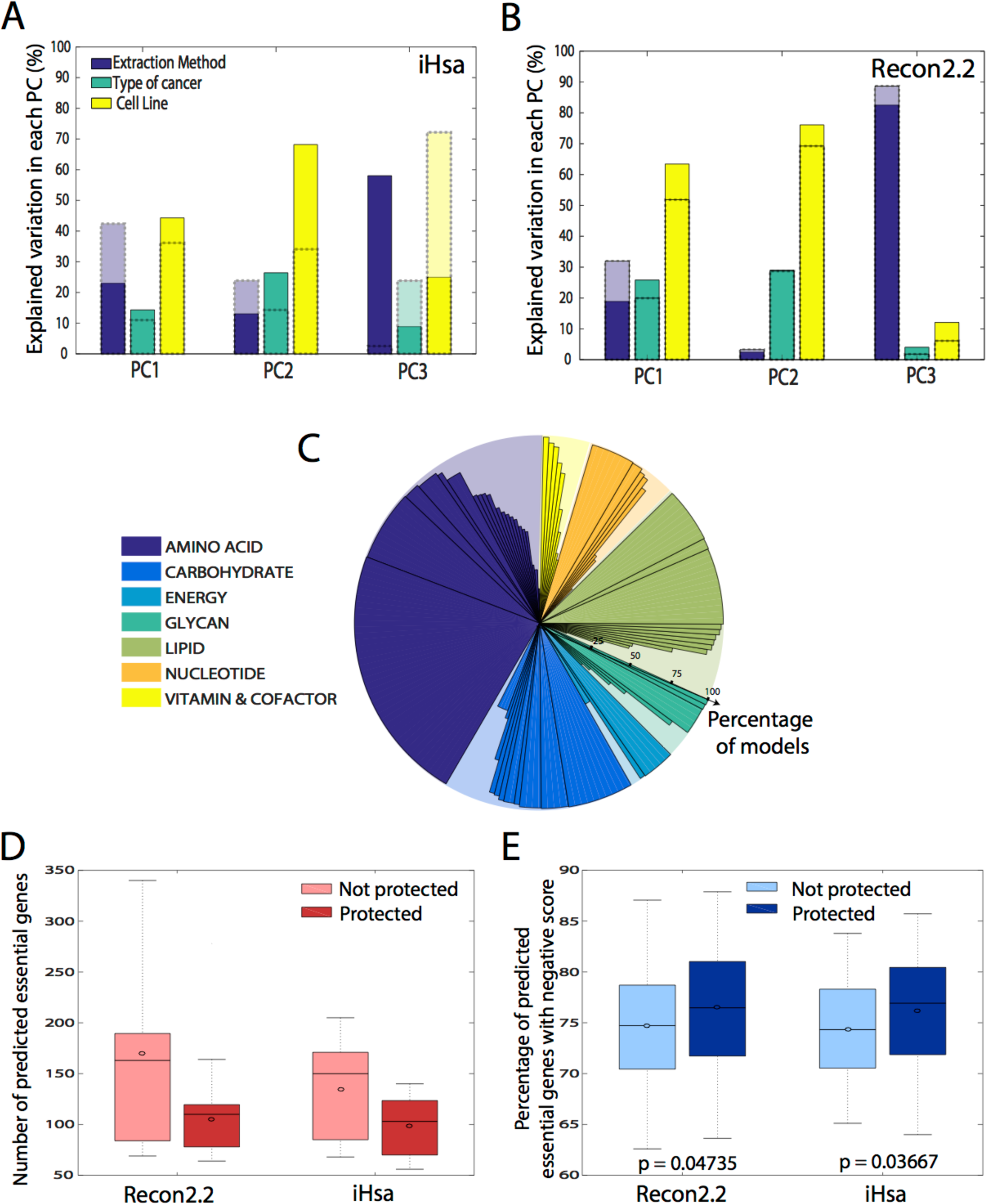
Protection of data-inferred tasks reduces the influence of extraction methods on model content. (A) and (B) After protection of tasks, the influence of the extraction method on the model content at the reaction level is decreased in the 3 first PCs for both reference models. These PCA analyses are based on the models generated using only the algorithms allowing a significant protection of the inferred metabolic task (fastCORE, mCADRE and MBA). (C) The implementation of the protectionist approach considerably increases the percentage of tasks present in all extracted models. (D) The protection of data-inferred metabolic tasks reduces the number of predicted essential genes (threshold of 90% WT growth) for both reference models. (E) However, task protection increases the percentage of predicted essential genes found in genome-wide CRISPR knock out screens (i.e., genes with a negative score). Significance of increase in percentage was evaluated using the 1-sided Wilcoxon rank-sum test, with the expectation that the protection should improve the prediction.

Beyond model content, we evaluated how the protectionist approach influenced model predictions. Thus we analyzed the influence of protecting inferred tasks on gene-essentiality predictions. We systematically deleted each gene in all generated models, and then used flux balance analysis to test models for normal or impaired growth. Gene deletions associated with impaired growth are considered as essential. We observe that task protection reduces the number of genes predicted to be essential for all thresholds considered (i.e., percentage of the maximum wild type growth rate), regardless of the reference model or the extraction method used (Figure 5D; Supplementary Figure 7). We further evaluated the accuracy of essentiality predictions by comparing these to CRISPR-Cas9 loss-of-function screens for 20 cell lines (Doench *et al*, 2016; Meyers *et al*, 2017; Aguirre *et al*, 2016). In these screens, essential genes are identified based on gene scores attributed using single guide RNA (sgRNA) abundance for each knockout before and after growth selection. Gene scores that are more negative have a higher probability of being essential. Therefore, the agreement between model predictions and the CRISPR screen data can be quantified as the percentage of predicted essential genes that have a negative gene score (Tobalina *et al*, 2016). Furthermore, the significance of the improvement gained from protecting data-inferred metabolic tasks can be computed using a 1-tailed Wilcoxon test. Consistent with previous reports (Gatto *et al*, 2015; Opdam *et al*, 2017), we found that the models, without protecting metabolic tasks, correctly predicted many essential genes. However, the protectionist approach provided an improvement to gene-essentiality predictions (Figure 5E; Supplementary Figure 8). The task protection, while reducing the number of predicted essential genes, increased the proportion of true positives.

## Discussion

Here we generated hundreds of models for 44 cell lines from the NCI-60 panel using multiple MEMs and two reference GeMs (Recon2.2 and iHsa) using standard approaches or by protecting metabolic tasks that have been directly inferred from transcriptomic data. We presented a comparative analysis of these two sets of models. As previously observed, the analysis of the first set of extracted models (i.e., models generated without protecting metabolic functions) indicated that the choice of model extraction algorithm significantly influenced the model content at the reaction level (Opdam *et al*, 2017; Ferreira *et al*, 2017; Robaina Estévez & Nikoloski, 2014). This leads to considerable variability in context-specific model content, which dwarfed the biological variability across cell lines, otherwise seen in their transcriptomes.

We provided here a curated list of 210 tasks that were used to compare the functionalities of the extracted models. The evaluation of metabolic tasks has emerged as a valuable practice in metabolic modeling studies (Agren et al., 2014; Blais et al., 2017; Bordbar et al., 2012; Chubukov, et al., 2014; Correia & Rocha, 2015; Duarte et al., 2007; Thiele et al., 2013; Uhlen et al., 2015). Such an approach allows one to evaluate the capacity of models to achieve specific modeling goals by capturing known metabolic features. Here we also demonstrated that the approach allows one to objectively compare models that may not share the same structure, such as different reference network reconstructions or models that have been extracted using different methods or parameters. We demonstrated that the selection of a reference model can significantly impact the resulting metabolic functions captured by extracted models, thus possibly impacting the results and interpretations from modeling studies. Indeed, the comparison of the functions of models extracted from both Recon2.2 and iHsa demonstrated the non-negligible influence of these reference models. We found this is principally due to differences in the GPR annotations in both GeMs. However, these differences in GPR annotations do not considerably influence the inference of metabolic tasks from transcriptomic data. The functional similarity across cell lines captured using data-inferred metabolic tasks is highly consistent between both reference models (Supplementary Figure 9). While community initiatives to standardize the formal representation of GeMs will facilitate cross comparison between numerous diverse existing GeMs (Lieven *et al*, 2018), these results highlight the potential of using the inference of functionalities directly from the transcriptome as a way to increase the consensus between extraction methods and reference models.

One challenge in the evaluation of metabolic models is the difficulty of comprehensively defining metabolic functions from a manual search of the literature. Thus, another strength of our approach is that it decreases the need for *a priori* knowledge or assumptions of the metabolic functions that should be included when building a cell or tissue specific model. Therefore, this list of metabolic tasks provides a framework for modelers to develop more physiologically accurate models by inferring the activity of metabolic tasks directly from omics data. Thus, key reactions that need to be included in a model can be protected, without requiring one to know what the cell does. However, the resulting models should still be curated to evaluate expected functionalities, such as for example auxotrophies.

Our protectionist approach can be implemented with diverse model extraction algorithms since it only requires the algorithms to prevent the removal of active metabolic tasks during the extraction process. However, some algorithms will require modifications to ensure the protection of all reactions related to a task. Current implementations of the GIMME-like and iMAT-like families do not favor this type of protection. By minimizing flux through reactions associated with low gene expression, GIMME-like extraction methods may remove low expression reactions one would want to retain for a validated metabolic task if there are high expression reactions that allow for growth. The iMAT-like methods are similar as they rely on finding an optimal trade-off between removing reactions associated with low gene expression, and keeping reactions whose genes/enzymes are highly expressed. Thus, modified implementations of these algorithms will be needed to allow the protection of reactions based on experimental observations. Finally, this approach can also be extended to any type of network complexity reduction that have been developed in the metabolic modeling field, such as the MILP-based approaches developed to tailor models based on exometabolomic data (Erdrich et al., 2015; Röhl & Bockmayr, 2017).

In our work, we also demonstrated that the models built with the protectionist approach are able to better capture cell-type specific metabolism and accurately predict many essential metabolic genes. Thus, these models may be invaluable for drug development strategies. The emergence of experimental techniques to assess the genetic vulnerabilities of a cell (e.g., CRISPR-cas, RNAi) allows researchers to identify sets of genes that should be essential for growth maintenance. These essential genes can further be used to evaluate the capacity of models to represent the interdependence between down-regulation of a gene and the concomitant impairment of growth. Thus, models can be used for interpreting the mechanisms underlying metabolic vulnerabilities that may be invaluable for new drug discoveries.

Finally, the list of tasks presented in this study was constructed based on existing repositories. However, a community effort could be undertaken to extend the scope and the definition of these metabolic functions. This would facilitate the description of genome-scale metabolic reconstructions as more than a network of reactions but rather as an interconnected map of cellular functions. This would be invaluable for the development of algorithms using more relevant biological information and facilitate more comprehensive and accurate descriptions of metabolic adaptations that occur in cells facing a change of context.

## Conclusion

Context-specific extraction methods are powerful approaches that provide insights in the metabolic state of a cell in specific environments. However, the underlying assumptions used to tailor the GeM based on omics data vary across algorithms, with the consequence that drastically different models can be obtained based on the same data. The poor consensus in generated models may limit the use of context-specific methods for data-driven hypotheses. The definition of metabolic tasks can help with these concerns. Our curated list of tasks and computational framework will allow users to infer metabolic functions directly from transcriptomic data using the whole genome-scale model, and drive the development of improved context specific models. Such models will pave the way toward a better consensus between existing context-specific extraction algorithms, and facilitate the application of models for novel biomedical and engineering applications.

## Methods

### Preprocessing of gene expression data

RNA-Seq data for the 44 cell lines from the NCI-60 panel were downloaded from CellMiner (Reinhold *et al*, 2012). We processed the gene expression data to attribute a gene activity score for each gene and define which genes are active in each cell line. A gene is defined as active in a sample if its expression value is above a threshold defined for this gene within the dataset considered. The threshold of a gene is defined by the mean value of its expression over all the samples coming from the same dataset with exceptions that the threshold needs to be higher or equal the 25^th^ percentile of the overall gene expression value distribution and lower or equal to the 75^th^ percentile. The gene score is computed as follows:

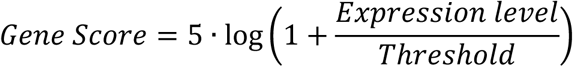

These gene scores are mapped to the models by parsing the GPR rules associated with each reaction. The gene score for each reaction is selected by taking the *minimum* expression value amongst all the genes associated to an enzyme complex (AND rule) and the *maximum* expression value amongst all the genes associated to an isozyme (OR rule) (Jensen *et al*, 2011). Note that we have recently benchmarked the influence of preprocessing methods on the definition of the set of active genes and observed that this parameter combination presented the best performance (Richelle *et al*, 2018).

### Implementation of the MEMs

Model extraction methods (MEMs) employ diverse algorithms to extract cell line- or tissue-specific models from a GeM. The MEMs we have considered can be categorized into three families: “GIMME-like” (i.e., GIMME), “iMAT-like” (i.e., iMAT and INIT) and “MBA-like” (i.e., MBA, FASTCORE, and mCADRE), as proposed previously (Robaina Estévez & Nikoloski, 2014). The GIMME-like family minimizes flux through reactions associated with low gene expression. The iMAT-like family finds an optimal trade-off between removing reactions associated with low gene expression, and keeping reactions whose genes/enzymes are highly expressed. In the MBA-like family, the algorithms use sets of core reactions that should be retained and active, while removing other reactions if possible. All the algorithms used in this study have been implemented using the function *createTissueSpecificModel* available in the COBRA Toolbox 3.0 (Heirendt *et al*, 2017). We describe below the list of required parameters needed to run the different methods, all optional parameters have been kept to their default setting.

#### FASTCORE

(Vlassis *et al*, 2013) **-** The core reactions set (*options.core*) is determined by all the reactions associated to a gene score superior to 5log(2). Note that the biomass reaction was added to the core reactions sets.

#### GIMME

(Becker & Palsson, 2008) **-** The implementation of GIMME requires two parameters: the gene scores (*options.expressionRxns*) and a threshold value, the reactions associated with a gene score value below this threshold will be minimized (*options.threshold* = 5log(2)). Note that we manually attributed a gene score of 10log(2) to the biomass reaction to ensure its inclusion.

#### IiMAT

(Shlomi et al., 2008; Zur et al., 2010) **-** Three parameters need to be provided to run iMAT: the gene scores (*options.expressionRxns*), a lower threshold value (reactions with gene score below this value are considered as “non-expressed”) and a upper threshold value (reactions with gene score above this value are considered as “expressed”). To simplify the comparison across algorithms, we set both thresholds to the same value: *options.threshold_lb= options.threshold_ub* = 5log(2), as done in a previous benchmarking study (Opdam *et al*, 2017). Note that we manually attributed a gene score of 10log(2) to the biomass reaction to ensure its inclusion.

#### INIT

(Agren *et al*, 2012) **-** The implementation of INIT requires attributing positive weights (*options.weights*) to each reaction with high expression and negative weights for the ones with low expression. All the reactions associated with a gene score below 5log(2) have been assigned a weight of −8 while the weights of remaining reactions were defined as the ratio between the gene score for each reaction and 5log(2). The weight associated with the biomass reaction was put to the maximum of obtained reaction weights.

#### MBA

(Jerby *et al*, 2010) **-** The implementation of MBA requires the definition of two set of reactions: high confidence (*options.high_set*) due to their expression and others with medium confidence (*options.medium_set*). The set of reactions with high confidence is defined as reactions with a gene score above the 75^th^ percentile of the distribution of all gene scores and the medium confidence set by all the reactions presenting score above 5log(2) and below the 75^th^ percentile of the distribution of all gene scores. Note that the biomass reaction has been manually added to the high confidence set of reactions.

#### mCADRE

(Wang *et al*, 2012a) **-** The implementation of mCADRE requires a score quantifying how often a gene is expressed across samples (*options.ubiquityScore*) and a literature-based evidence score (*options.confidenceScores*). Since the confidence score identification used in the original paper is difficult to transpose in this study, we did not define the confidence score as preformed in the tutorial presenting the implementation of mCADRE in COBRA Toolbox 3.0 (Heirendt *et al*, 2017). Furthermore, as the gene scores are computed based on the knowledge of the gene expression of a gene across all samples, we used the gene scores as ubiquity scores.

### Curation of metabolic tasks

The curation has been done by first taking the union of previously published lists of metabolic tasks (Thiele *et al*, 2013; Blais *et al*, 2017). We removed duplicated tasks and lumped tasks that rely on the description of similar metabolic functions. Each remaining task without strong biological evidence was removed. We also created 9 new tasks that were essential for the acquisition of already described metabolic functions (i.e., intermediate biosynthetic steps for the acquisition of other tasks). Doing so, we obtained a collection of 210 tasks associated with 7 systems (energy, nucleotide, carbohydrates, amino acid, lipid, vitamin & cofactor and glycan metabolism). For each task, we provided its original source (Recon and/or iHsa) and comments on the biological evidence of this metabolic function (Supplementary Table 4).

### A unified framework for computing metabolic tasks as a model benchmarking tool

In its original version, Thiele and coworkers (2013) define a “*metabolic task as a nonzero flux through a reaction or through a pathway leading to the production of a metabolite B from a metabolite A. The metabolic capacity of the network was demonstrated by testing nonzero flux values for these metabolic tasks. For each of the simulations, a steady-state flux distribution was calculated. Each metabolic task was optimized individually by choosing the corresponding reaction in the model, if present, as objective function and maximized the flux through the reaction*”. In parallel, Agren and coworkers presented an alternative framework to compute the metabolic tasks present in a model within their RAVEN toolbox (Agren *et al*, 2013). They defined a metabolic task through a list of inputs and outputs for which the pseudo-stationary assumption will be relaxed following a magnitude imposed by the user and assumed that a task successfully passes if the variation imposed to the inputs leads to the imposed variation of the outputs.

We also propose to define a metabolic task as the capacity of producing a defined list of output products when only a defined list of input substrates is available. However, we modified the way to implement it from the RAVEN toolbox. Instead of relying on the relaxation of the steady-state assumption, we take an approach more similar to that proposed by Thiele et al (2013) by imposing constraints only at the flux level. Therefore, a model successfully passes a task if the associated LP problem is still solvable when the sole exchange reactions allowed carrying flux in the model are temporary sink reactions associated with each of the inputs and outputs listed in the task. This framework allows the use of known stoichiometry to fix the ratio between the fluxes of the sink reactions associated with each input and output of the task. We implemented the code to compute the tasks in Matlab, and the code, *checkMetabolicTasks*, has been contributed to the COBRA Toolbox3.0 (Heirendt *et al*, 2017).

### Validation on existing animal genome-scale models

We tested the list of tasks using published genome-scale models of human (Swainston *et al*, 2016a; Thiele *et al*, 2013; Duarte *et al*, 2007; Quek *et al*, 2014; Blais *et al*, 2017), Chinese hamster (Hefzi *et al*, 2016), rat (Blais *et al*, 2017) and mouse (Sigurdsson *et al*, 2010) cells (Figure 2D, Supplementary Table 5). All models successfully pass more than 90% of the tasks. For each failed task, we provided a reason of the failure (i.e. definition of the missing reaction to successfully pass the task) (Supplementary Table 5). As the definition of the metabolic tasks depends on the provision of the exact name of the metabolites in each model, we also provide a table of nomenclature compatibility between the different genome-scale models tested (Supplementary Table 6).

### Inference of metabolic tasks from transcriptomic data

We developed a computational framework for attributing a score to each metabolic task in order to extend the application of the concept beyond the model benchmarking scope. If a task successfully passes in a model, one can compute the list of reactions associated with this task and, in doing so, access the list of genes that may contribute to the acquisition of this metabolic function based on the GPR rules. To this end, we used the parsimonious Flux Balance Analysis (pFBA) algorithm to define the set of reactions and associated genes required to pass a task within a specified model (Lewis *et al*, 2010). Thanks to the availability of this information, metabolic functions can now be directly assessed from transcriptomic data. The proposed computation of a metabolic score relies first on the preprocessing of the available transcriptomic data and the attribution of a gene activity score for each gene (see associated Methods section). We further used the GPR rules associated with each reaction required for a task to decide which gene will be the main determinant of the enzyme abundance associated with this reaction and attribute the corresponding gene activity level. Therefore, each reaction involved in a task is associated with a reaction activity level (RAL) that corresponds to the preprocessed gene expression value of the gene selected as the main determinant for this reaction. Finally, the metabolic score can be computed as the mean of the activity level of each reaction:

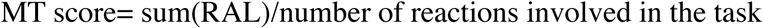

Doing so, a metabolic task will be considered as active if its MT score has a value greater than 5log(2). The list of active metabolic tasks for each of the 44 cell lines from the NCI-60 panel is available in Supplementary Table 7.

### Protection of data-inferred task during extraction process

We used the list of active metabolic tasks (Supplementary Table 7) to determine the set of reactions that should be protected during the extraction process for each of the 44 cell lines. The protectionist approach has been implemented for each extraction method by using the same set of parameters as previously described with the following modification:

#### FASTCORE

The set of reactions associated with the metabolic tasks defined as active based on the transcriptomic data has been manually added to the core reactions set (*options.core*).

#### GIMME & iMAT

A gene score of 10log(2) (*options.expressionRxns*) has been attributed to all the reactions associated to the metabolic tasks defined as active based on the transcriptomic data.

#### INIT

The weights (*options.weights*) for all reactions associated with the metabolic tasks defined as active based on the transcriptomic data were put to the maximum of obtained reaction weights.

#### MBA

The reactions associated with the metabolic tasks defined as active based on the transcriptomic data have been manually added to the high confidence set of reactions.

#### mCADRE

A ubiquity score (*options.ubiquityScore*) of 1 has been attributed to all the reactions associated to the metabolic tasks defined as active based on the transcriptomic data.

### Principal component analysis

For the reaction PCAs, a binary matrix is constructed in which each row represents an extracted model and each column represents a reaction, with each element representing the presence (1) or absence (0) of a reaction in a model. Reactions in all or no models were removed from the matrix. Similarly for the metabolic function PCA, the matrix had each row as an extracted model and each column as a metabolic task, with each element in the matrix representing if the task is present (1) or absent (0) in a model. For the PCAs, the matrix was centered to have zero mean within each row. PCA was done on this matrix. The variance explained by the different factors (MEM, cancer type and cell line) within each of the principal components is calculated as follows. Within one factor, the maximum Pearson correlation coefficient (R) of the component scores and categories is calculated across all possible orderings of the categories. Reported is the R^2^ scaled to percentages. The same procedure was used to perform the PCA on the model functionalities except that the binary matrix of reactions was replaced by the binary matrix representing the list of metabolic tasks that are successfully passed in each extracted model. The attributes of all extracted models (number of reactions and metabolites, number of successfully passed tasks and predicted growth rate) are available in Supplementary Table 8 and the results of the extracted model benchmarking using the list of metabolic tasks is available in Supplementary Table 9.

### Predictions of Gene-Essentiality

To predict gene-essentiality, FBA was used to optimize biomass production following the removal of each reaction in the cell line-specific models that would be affected by gene removal based on the GPRs. The function used to perform this deletion analysis is available in COBRA Toolbox 3.0, *singleGeneDeletion.m* (Heirendt *et al*, 2017). To test these essentiality predictions of the models against experimental data, we downloaded CRISPR-Cas9 loss-of-function screens data for 20 NCI-60 cell lines from depmap.org (Doench *et al*, 2016; Meyers *et al*, 2017; Aguirre *et al*, 2016). In these screens, essential genes are identified based on genes scores attributed using single guide RNA (sgRNA) abundance for each knockout before and after growth selection. A more negative gene score suggests a higher probability that the gene is essential. Therefore, the agreement between prediction and data can be analyzed by using the percentage of predicted essential genes that have a negative gene score (Tobalina *et al*, 2016). A 1-tailed Wilcoxon rank sum test was used to test whether the percentage of predicted essential genes of the model extracted using the protectionist approach were significantly higher than the ones without protection. The results of the gene deletion study and prediction against CRISPR-Cas9 loss-of-function screens are available in Supplementary Table 10.

## Acknowledgements

This work was supported by generous funding from the Novo Nordisk Foundation provided to the Center for Biosustainability at the Technical University of Denmark (NNF10CC1016517), and from NIGMS (R35 GM119850). We also thank the Lilly Innovation Fellowship program for generous funding provided to AR that enabled this work.

## Author contribution

AR, and NEL designed the study, conducted the analyses, and wrote the paper. AC and CK analysed data. All authors have read and approved the work.

